# Optogenetics reveals roles for supporting cells in force transmission to and from outer hair cells in the mouse cochlea

**DOI:** 10.1101/2023.06.26.546543

**Authors:** Victoria A. Lukashkina, Snezana Levic, Patricio Simões, Zhenhang Xu, Yuju Li, Trevor Haugen, Jian Zuo, Andrei N. Lukashin, Ian J. Russell

**Author notes:** Sussex Neuroscience, School of Life Sciences, University of Sussex, Brighton BN1 9QG, UK. **Funding:** This work was funded by United Kingdom Medical Research Council Grant MR/W028956/1.

## Abstract

Cochlear outer hair cells (OHCs), acting as bidirectional cellular mechanoelectrical-transducers, generate, receive, and exchange forces with other major elements of the cochlear partition, including inner hair cells (IHCs). Force exchange is mediated via a supporting cell scaffold, including Deiters’ (DC) and outer pillar cells (OPC), to enable the sensitivity and exquisite frequency selectivity of the mammalian cochlea. We conditionally expressed a hyperpolarizing halorhodopsin (HOP), a light-gated inward chloride ion pump in DCs and OPCs. We measured extracellular receptor potentials (ERPs) and their DC component (ERPDC) from the Cortilymph (CL) of HOP expressing mice and compared the responses with similar potentials from littermates without HOP expression. Compound action potentials (CAP) were measured as an indication of IHC activity. HOP laser activation suppressed cochlear amplification through changing timing of its feedback, altered basilar membrane (BM) responses to tones at all measured levels and frequencies, and reduced IHC excitation. Our HOP activation results here complement previous channelrhodopsin activation studies in exploiting optogenetics to measure and understand the roles of DCs and OPCs *in vivo* in controlling the mechanical and electrical responses of OHCs to sound and their contribution to timed and directed electromechanical feedback to the mammalian cochlea.

**SIGNIFICANCE STATEMENT:** Outer hair cells provide electromechanical feedback to the organ of Corti, mediated via a cellular scaffold of Deiters’ and outer pillar cells, that enables the sensitivity and fine frequency tuning of the cochlea. The role of this scaffold was explored by expressing the halorhodopsin HOP in Deiters’ and pillar cells which, when illuminated, hyperpolarized them. HOP activation suppressed cochlear amplification through altering the timing of outer hair cell forces to the Organ of Corti, altered basilar membrane responses to tones, including those at levels and frequencies not subject to amplification, and reduced neural excitation. The findings implicated roles for supporting cells in mediating force transmission to and from outer hair cells along all axes of the organ of Corti.

## INTRODUCTION

The ability of the cochlear outer hair cells (OHCs) to generate, receive and exchange forces with other major elements of the cochlear partition, including the tectorial membrane (TM), basilar membrane (BM) and sensory inner hair cells (IHCs) (Fig. 1A), enables the sensitivity and exquisite frequency selectivity of the mammalian cochlea. OHCs, acting as bidirectional cellular mechanoelectrical-electromechanical transducers and amplifiers, interact with the major noncellular elements of the cochlear partition and IHCs via a supporting cell scaffold, including Deiters’ cells (DCs) and outer pillar cells (OPCs) (Fig.1B). Our current understanding of the possible roles of the cellular scaffold in the transfer of OHC forces transversely, radially, and longitudinally in the cochlear partition is based largely on modelling studies and from *ex vivo* preparations rather than from *in vivo* measurements (Yoon et al., 2011; Nam, 2014; Sasmal and Grosh, 2019; Zhou et al., 2022).

**Figure 1.**
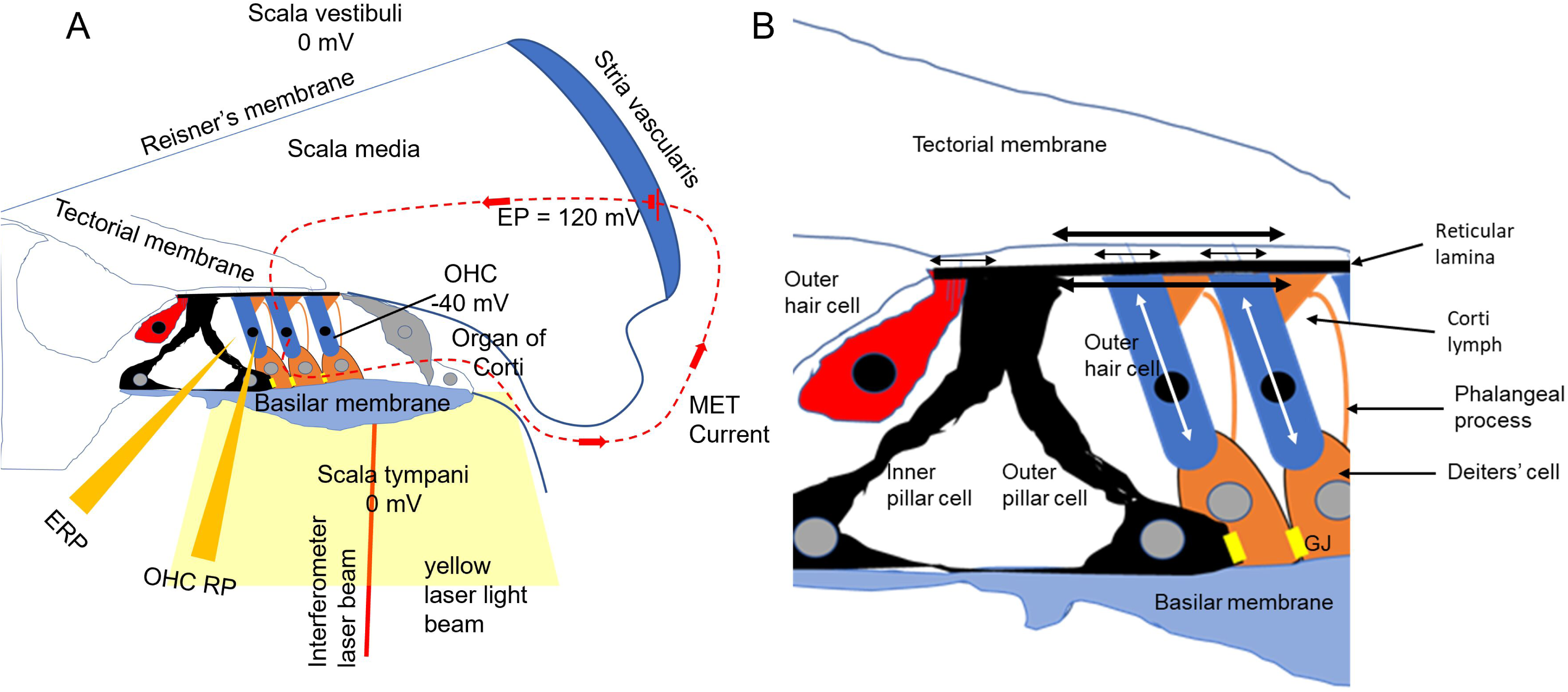
Schematics of cochlea. **A**. Schematic cross-section of cochlea, including OoC, with three fluid-filled chambers (scala tympani, scala media, and scala vestibuli) separated by the BM and Reisner’s membrane. The positive endocochlear potential (EP) in series with the negative OHC membrane potential provides the battery driving the K^+^ dominated, MET current pathway (dashed lines). Micropipettes were used to record extracellular potentials from the CL and intracellularly from OHCs. A red self-mixing interferometer laser beam was used to measure BM displacements. A yellow laser beam activated HOP expressed mainly in DCs. **B**. OHC MET current is modulated when the OHC MET conductance is altered by sound-induced OHC bundle displacement (small, horizontal double arrows) due to radial, shear displacements between the TM and the RL (large double arrows) caused by transverse sound-induced BM vibrations (large vertical double arrows). Transmembrane voltage changes caused by the modulated MET current trigger length and force changes in the OHCs (vertical white double arrows) with positive intracellular potentials causing OHCs to shorten.

OHCs exchange forces locally with other elements of the cochlear partition at their tonotopic frequency (characteristic frequency, CF) location in the organ of Corti (OoC) that is determined by the mechanical properties of the BM (Fig. 1), which is graded in increasing stiffness from the low-frequency apex to the high-frequency base of the cochlea (Robles and Ruggero, 2001). OHCs receive sound induced transverse forces via the vibrations of the BM and their supporting cell cages, which results in radial shear displacements between the reticular laminar and the overlying TM (Fig. 1B). The radial shear displaces the OHC hair bundles and modulates the open probability of the mechanoelectrical transduction (MET) conductances that are present at the tips of all but the tallest row of stereocilia of the hair bundles (reviewed by Fettiplace, 2017). OHCs generate forces as a consequence of the voltage differences generated across the OHC basolateral membranes due to the modulated flow of MET current (Fig. 1A), which controls the prestin based voltage dependent somatic motility (reviewed by Ashmore, 2008; Dallos, 2008; Robles and Ruggero, 2001; Levic et al., 2022). In vivo measurements (Nilsen and Russell, 1997; 2000) and models (Geisler, and Sang, 1995: Nobili and Mammano, 1996;) reveal that optimal timing of force delivery to the cochlear partition occurs when OHCs shorten during maximum BM velocity towards the scala media. Are there roles for the supporting cell cage in enabling the optimum timing of OHC feedback to the cochlear partition and in the transmission of forces between OHCs and other structures in the cochlear partition? To address this question, we used *in vivo* and *ex vivo* optogenetic techniques based on the conditional expression in DCs and OPCs of the halorhodopsin (HOP), which gates an inward chloride ion pump that reversibly hyperpolarizes cell membrane potentials when light-activated (Zhang et al., 2007). We measured extracellular receptor potentials (ERPs) and their DC component (ERPDC) from the Cortilymph (CL) immediately adjacent to the OHCs as close approximations of OHC intracellular responses (Kössl and Russell, 1992; Levic et al., 2022), and compared similar potentials recorded from the same cochlear frequency location from littermates without HOP expression. We measured local compound action potentials (CAP) as an indirect indication of IHC activity. We discovered that activation of HOP expressed in DCs altered the phase of the ERP, suppressed cochlear amplification, altered sound evoked responses of BM at all measured levels and frequencies, and suppressed IHC excitation.

## MATERIALS AND METHODS

The Animal Resource Facilities of Creighton University approved protocols on mouse breeding, husbandry and cochlear morphological analysis performed in this study. All *in vitro* and *in vivo* experiments performed at the University of Brighton complied with Home Office guidelines under the Animals (Scientific Procedures) Act of 1986 and were approved by the University of Brighton Animal Welfare and Ethical Review Body.

Mice were housed in a facility with a 12-h light/dark cycle and free access to food and water. All mice used here were purchased from The Jackson Lab: Fgfr3iCreER^T2^ (stock # 025809) and Rosa-CAG-LSL-eNpHR3.0-EYFP-WPRE or HOP-EYFP (Ai39; stock #014539) were crossed to create the Fgfr3-iCreER^T2^; HOP-EYFP mouse. Tamoxifen was injected intraperitoneally (i.p.) at 250 mg/kg at P21 mice to visualize the expression of HOP-EYFP in cochleae at P28. Mice were genotyped as described in protocols by The Jackson Lab.

To induce HOP-EYFP expression specifically in mature DCs and OPCs in the OoC, we used a previously characterized Fgfr3-iCreER^T2^ mouse line that displays ∼100% inducible Cre activity in cochlear DCs and OPCs when tamoxifen was injected at juvenile and adult ages (Cox et al., 2012, Walters et al., 2017). No HOP-EYFP was detected in DCs/PCs in Fgfr3-iCreER^T2-^; HOP-EYFP^+^ control mice at P28, whereas Fgfr3-iCreER^T2+^; HOP-EYFP^+^ experimental mice showed expression of HOP-EYFP fusion protein robustly in DCs and weakly in PCs within the OoC at P28 when induced with tamoxifen at P21. Both autofluorescent and immunofluorescent signals of EYFP were detected in the phalangeal processes of DCs and PCs of experimental mice. We will use HOP in brief for HOP-EYFP afterwards.

The cochlear samples were fixed in 4% paraformaldehyde for overnight at 4 °C (P28 cochleae). Tissues were washed in PBS and decalcified in 120mM EDTA (pH = 7.4) for 48 hrs before microdissection of the OoC. The samples were blocked and permeabilized in the blocking buffer (10% Fetal Bovine Serum serum and 0.2 % Triton X-100 in PBS) for 2 hrs at room temperature. They were then incubated at 4 °C overnight with primary antibody rabbit anti-Myosin VI (1:400 Proteus Bioscience) and mouse anti-green fluorescent protein (Invitrogen, cat. A11120). The tissues were incubated with 1:800 diluted secondary antibodies for 2 hrs at room temperature, washed with PBS, and incubated with Dapi (1:1000 Thermo Scientific) for 2 mins at room temperature, washed with PBS, and then mounted for imaging using Fluoromount-G (SouthernBiotech). All samples were imaged with an LSM700 confocal laser scanning image system (Carl Zeiss, Jena, Germany).

### In vitro methods

For physiological *in vitro* and *in vivo* experiments, tamoxifen was injected intraperitoneally at 250 mg/kg at days P12 and P13 to induce HOP expression in DCs and OPCs by P21 (Lukashkina et al., 2022). *In vitro* methods used were similar to Lukashkina et al., (2022). P17-P20 male and female mice were killed using cervical dislocation. The cochlea was dissected and isolated in an ice-cold solution containing (in mM): 135 NaCl, 5.8 KCl, 1.3 CaCl2, 0.9 MgCl2, 0.7 NaH2PO4, 5.6 D-glucose, 10 HEPES, 2 sodium pyruvate, pH 7.5 (adjusted with NaOH) and osmolarity 308 mOsm.

In order to investigate the DC electrical behavior under illumination within the syncytium, the apical coil of the OoC was immobilized with a nylon mesh fixed to a stainless-steel ring in the recording chamber and viewed using an upright microscope with Nomarski differential interface contrast optics (40x water immersion objectives, Axioskope). HOP, a halorhodopsin encoding a light-gated chloride channel (Schobert and Lanyi, 1982), was activated using 30mW 589 nm DPSS Yellow Laser system. To assess the cellular effects of light activation of HOP on DC electrical responses, membrane potentials were recorded using the same solution as for dissection with the addition of 100 μM carbenoxolone to block gap junctions. All recordings were made at room temperature (22°C). Patch pipettes were filled with an intracellular solution containing the following (in mM): 131 KGluconate, 3 MgCl_2_, 5 Na_2_ATP, 10 Na2-phosphocreatine, 1 EGTA, 10 HEPES, 0.3 Na_2_GTP, pH 7.4 and osmolarity 293 mOsm. All chemicals were obtained from Sigma Aldrich. The patch pipettes were fabricated with a dual-stage glass micropipette puller (Narishige PC-100) using borosilicate glass with outer diameter 1.5 mm (Sutter Instruments) and heat polished with a microforge (Narishige MF-900). Currents were amplified with an Axopatch 700B amplifier (Molecular Devices) and filtered at a frequency of 2-5 kHz through a low-pass Bessel filter. The data were digitized at 5-20 kHz using an analog-to-digital converter (Digidata 1500; Molecular Devices). The whole-cell current recordings were conducted using pCLAMP software (version 10, Molecular Devices).

### *In Vivo* Physiological Recordings

*In vivo* methods are similar to those described in Lukashkina et al (2022). Mice at 3-5 weeks of age were anaesthetized with ketamine (0.12 mg/g body weight i.p.) and xylazine (0.01 mg/g body weight i.p.) for nonsurgical procedures or with urethane (ethyl carbamate; 2 mg/g body weight i.p.) for surgical procedures at the University of Brighton. Mice were tracheotomized, and their core temperature was maintained at 38 °C. The auditory sensitivity of mice was assessed before surgery using distortion product otoacoustic emissions (DPOAE, see below) to ensure each mouse was sensitive throughout the 1-70 kHz range of the sound system and especially sensitive to tones in the 50 kHz – 60 kHz range to levels ≤ 30 dB SPL.

To measure BM displacements and the OoC cochlear microphonics (CM), a caudal opening was made in the ventro-lateral aspect of the right bulla to reveal the round window (RW) (Legan et al., 2000). Sound was delivered via a probe with its tip within 1 mm of the tympanic membrane and coupled to a closed acoustic system comprising two MicroTechGefell GmbH 1-inch MK102 microphones for delivering tones and a Bruel and Kjaer (www.Bksv.co.uk) 3135 0.25-inch microphone for monitoring sound pressure at the tympanum and DPOAE recording. The sound system was calibrated *in situ* for frequencies between 1 and 70 kHz and known sound-pressure levels were expressed in dB SPL with reference to 2×10^−5^ Pa. Tone pulses with rise/fall times of 1 ms were synthesized by a Data Translation 3010 (Data Translation, Marlboro, MA) data acquisition board, attenuated, and used for sound-system calibration and the measurement of electrical and acoustical cochlear responses. To measure DPOAEs, primary tones were set to generate 2f1−f2 distortion products at frequencies between 1 and 50 kHz. DPOAEs were measured for f1 levels from 10 to 80 dB SPL, with the levels of the f2 tone set 10 dB below that of the f1 tone. DPOAE threshold curves represented the level of the f2 tone that produced a 2f1− f2 DPOAE with a level of 0 dB SPL when the f2/f1 frequency ratio was 1.23. System distortion during DPOAE measurements was 80 dB below the primary-tone levels.

Extracellular receptor potentials (ERPs) were recorded close to OHCs from the fluid spaces of the OoC from wild-type (WT) and HOP mice OHCs and intracellular receptor potentials were recorded from presumed OHCs from WT mice (Fig. 1A) using glass pipettes (20 MΩ - 80 MΩ when filled with 3M KCl) pulled from 1 mm diameter thin-walled quartz glass tubing on a Sutter P-2000 micropipette puller (Sutter Instrument Novato, CA 94949, USA). Signals were amplified with a recording bandwidth of DC - 100 kHz using a custom preamplifier (Brighton lab, James Hartley). The voltage signals were not capacitance compensated. Presumed DCs with very negative resting membrane potentials (-112.5 ± 8.3 mV, n = 18) were encountered by advancing microelectrodes through the RW membrane and the BM in the close vicinity of OHCs in the 50 kHz - 60 kHz region of the basal turn of the cochlea. Further advance resulted in encountering the OoC fluid spaces (i.e. CL) that are adjacent to the OHCs, with zero potentials, and scala media with positive EP (+114.3± 3.7 mV, n=11).

Tone-evoked BM displacements were measured by focusing the beam of a self-mixing, displacement sensitive, laser-diode interferometer (Lukashkin et al., 2005) through the RW membrane to form a 20-μm spot on the center of the BM in the 50-60 kHz region of the cochlea (Fig. 1B). To take rapid snapshots of the BM responses without many averages, the following algorithm was employed: The response amplitude of the interferometer is the largest when its operating point is situated in quadrature (Lukashkin et al., 2005). During recordings, the operating point fluctuated due to physiological noises in the preparation, and we constantly tracked for a response in, or very close to, quadrature, i.e. for the largest response. We balanced the demand for rapid data acquisition with the need for sensitive measurement by spotting the maximum response in 10-15 presentations of the same tone burst. The largest response amplitude recorded was the one closest to the quadrature response. This allowed us to complete recording of the entire level functions (dependence of the BM movement on stimulation level) within 15-20 seconds. The interferometer was calibrated at each measurement location by vibrating the piezo stack on which it was mounted over a known range of displacements. BM measurements were checked continuously for changes in the sensitivity of the measurement (due to changes in alignment or to fluid on the RW) and for changes in the condition of the preparation. If thresholds of the latter changed by more than 5-10 dB, the measurements were terminated. Tone pulses with rise/fall times of 1 ms were used for BM measurements. Stimulus delivery to the sound system and interferometer for calibration and processing of signals from the microphone amplifiers, microelectrode recording amplifiers, and interferometer were controlled by a DT3010/32 (Data Translation, Marlboro, MA) board by a PC running Matlab (The MathWorks, Natick, MA) at a sampling rate of 250 kHz. The output signal of the interferometer was processed using a digital phase-locking algorithm, and instantaneous amplitude and phase of the wave were recorded.

To activate HOP, the BM was illuminated by a 30 mW 589nm DPSS Yellow Laser System (Dragon Lasers, Changchun Jilin, China) coupled to a fiber-optic cable (Thorlabs, M63L01, 105μm, 0.22NA). The tip was positioned with a micromanipulator to be 0.2 mm from the surface of the RW membrane, where it cast a ∼200 µm diameter circle of illumination on the BM, centered on either the micropipette or the BM displacement measurement beam (Figure 1A). The distance of 0.2 mm from the RW was determined during the calibration of the beam by casting a 200 µm diameter circle of illumination on the surface of the photo diode sensor of a power meter (Thorlabs, PM16-130). The laser power at the level of the OHCs was computed according to Wu et al. (2016), who used analysis and estimates provided by Zhang et al. (2007) and Aravanis et al. (2007). Laser on-off was controlled through transistor-transistor logic (TTL).

Measurements were made without knowledge of genotype. Less than 5% of all measurements were terminated because the physiological state of the preparation changed during measurements, in which case data from the sample was excluded.

### Experimental design and statistical analyses

HOP^+/-^ experimental mice were crossed to generate +/+, +/- and -/- genotypes. Male and female mice were studied in approximately equal proportions. No phenotypic differences were observed between males and females. Physiological tests were performed on +/+ and -/- littermates to minimize any influence of age, environment or genetic background. Tests were performed on 3 – 5-week-old mice to reduce the possibility of progressive loss of high frequency responses, which is common in many mouse strains. Tests were performed on all mice in a litter without knowledge of genotype. The genotypes were determined after the experiments. For statistical analysis of physiological experiments, data were compared for at least 5 +/+ and 5 HOP-/- mice, obtained from recordings of 2 or 3 complete litters. For analysis of BM and electrophysiological measurements from presumed DCs and fluid spaces of the OoC, 18 +/+ and 10 -/- HOP mice in total were tested. Data were analyzed as mean ± standard deviation (S.D.) and plotted using Fig P (www.figpsoft.com) or Origin (www.originlab.com) software. Statistical tests were performed with GraphPad Prism (https://www.graphpad.com/quickcalcs) and comparisons made using unpaired *t* tests for unequal variances unless otherwise noted. P values are noted as absolute values with *t* values and degrees of freedom (df).

In summary, all measurements were performed blind. Measurements were made from each animal in a litter and data were analyzed at the end of each set of measurements. When all measurements had been made from a particular litter, the tissue was genotyped. Phenotypic differences between the WT, heterozygous (HOP+/-) and homozygous (HOP+/+) mice were very strong. Thus, only sufficient numbers of measurements were made to obtain statistically significant differences. Experiments were terminated (<5% of all measurements) if the physiological state of the preparation changed during a measurement and data from the measurement was excluded.

### Software / code

For physiological recordings, data acquisition and data analysis were performed using a PC with programs written in MATLAB (The MathWorks, Natick, MA). The programs are available upon request from authors ANL and IJR. Please note that the programs were written to communicate with specific hardware (Data Translation 3010 board and custom-made USB-controlled attenuators) and will need modification if used with different hardware.

## RESULTS

### Specific inducible expression of HOP in cochlear supporting cells

To induce HOP-EYFP expression specifically in mature DCs and PCs in the OoC, we used a previously characterized Fgfr3-iCreER^T2^ mouse line that displays ∼100% inducible Cre activity in cochlear DCs and PCs when tamoxifen was injected at juvenile and adult ages (Cox et al., 2012, Walters et al., 2017). Similar to our previous studies on ChR2-tdTomato or COP (Lukashkina et al., 2022), no HOP-EYFP was detected in DCs/PCs in Fgfr3-iCreER^T2-^; HOP-EYFP^+^ control mice, whereas Fgfr3-iCreER^T2+^; HOP-EYFP^+^ experimental mice showed expression of HOP-EYFP fusion protein specifically in both DCs and PCs within the OoC at P28 post-tamoxifen induction. No expression of HOP-EYFP was detected in stria vascularis or any other cochlear structures of Fgfr3-iCreER^T2+^; HOP-EYFP+ experimental mice at P28 post-tamoxifen induction. These results on HOP-specific expression in DCs and PCs in the cochlea are consistent with previous studies (Hayashi et al., 2010; Cox et al., 2012; Walters et al., 2017). We will use HOP in brief for HOP-EYFP afterwards.

### Light hyperpolarizes HOP expressing Deiters’ cells

To examine the effects of light-activation of HOP expressed in DCs, we used whole cell current clamp configuration to make measurements from single DCs in *ex-vivo* flat mounted preparations of the OoC. The preparation was bathed in a solution (see methods) containing 100 μM carbenoxolone that blocked gap junctions. We measured voltage responses to current injections from -80pA to +80 pA in 10pA increments (Fig. 2A). Exposure to 1 second laser illumination caused V-I curves of DCs that expressed HOP to be shifted to the right in proportion to the power density of the illumination (Fig. 2A). DCs that did not express HOP were insensitive to laser illumination (Fig. 2A). In Fig. 2B, the voltage changes due to current injection without HOP activation have been subtracted from voltage changes with HOP activation at given injected currents to reveal the voltage changes due to HOP activation. DCs were hyperpolarized by HOP activation. Hyperpolarization was increased with increased DC membrane potential negativity and with increased illumination power density (Fig. 2B). Inset illustrates responses to a period of laser illumination (red line) from control and HOP expressing DCs. The responses display a fast onset and do not adapt to the laser illumination.

**Figure 2.**
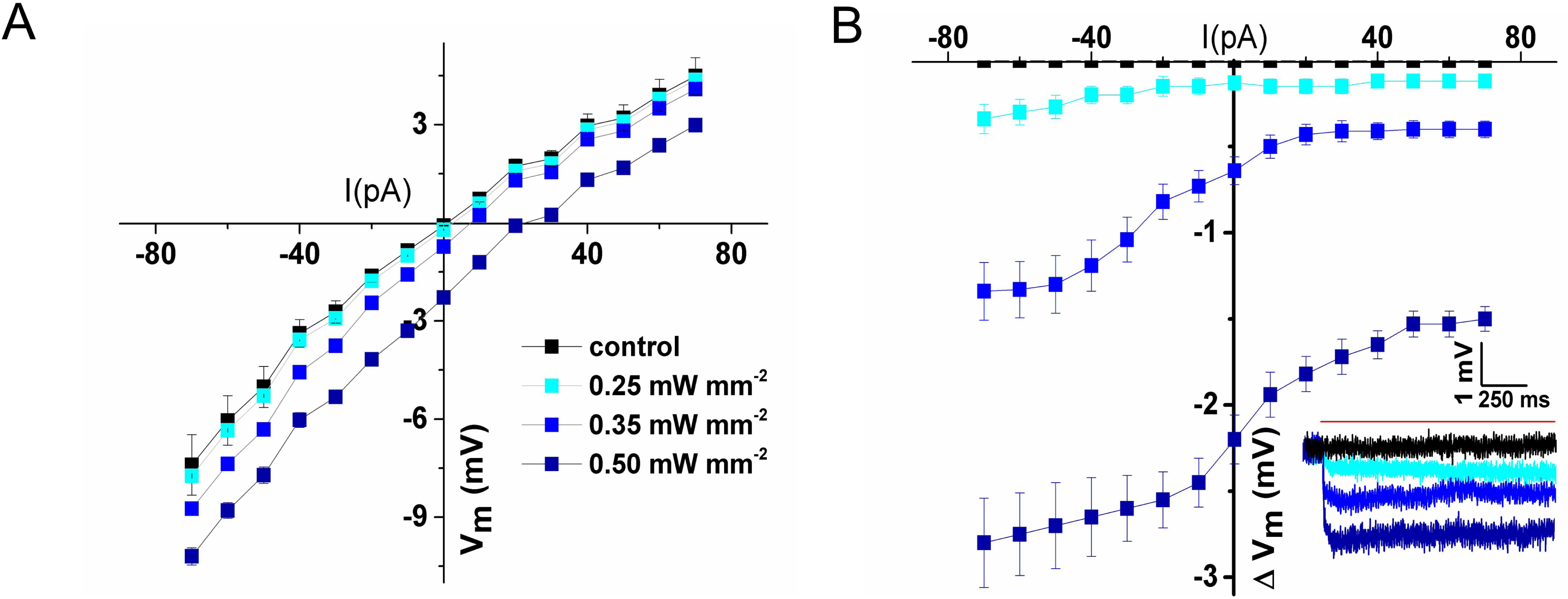
Illumination hyperpolarizes HOP expressing Deiters’ cells. **A**. Relationship between injected current (I) and change in membrane potential (V_m_) of HOP expressing (blue symbols) and non-HOP-expressing (control, back symbols) DCs during different levels (power densities) of laser illumination. **B**. The same data as shown in **A** but the voltage changes due to current injection alone have been subtracted from voltage change elicited due to current injection during HOP-activating laser illumination to reveal the voltage changes due to HOP activation. The average potential changes ± S.D. measured from a resting membrane potential of -50 mV for laser power densities 0.25, 0.35 and 0.50 mW mm^-2^ shown in B, are (in mV) -0.34±0.09, -1.34 ±0.17, and -2.81±0.26, respectively (n = 6). Inset, examples of single traces from whole-cell current-clamp recordings (resting membrane potential of -50 mV) showing responses to light-elicited hyperpolarization in response to a period of HOP laser activation (red line) at the power densities indicated. Current steps: 70 pA. Laser illumination 200 µm diameter, 589 nm.

### HOP activation reduces the endocochlear potential (EP)

The EP (+117 ± 4 mV, n = 5) measured from the scala media of mice that express HOP in DCs and OPCs, but not in littermates that did not express HOP (EP: 118 ± 4 mV, n = 7), is reduced when the HOP is activated by laser illumination (Fig. 3). The initial decrease of ∼ 2 mV (2.3 ±0.5 mV, n = 5) takes place in about 500 ms with a further decline to ∼2.5 mV (2.7 ±0.3 mV, n = 5) after about 4 s of light exposure. If the period of light exposure is < 4s, the EP returns to its original level after turning off the laser light. For longer periods of illumination, the EP continues to change with further decreases and increases (Fig. 3). For this reason, laser illumination of the BM was confined to periods of < 4 s. Thus, light activation of HOP expressed in DCs causes hyperpolarization in DCs (Fig. 3) and a small reduction in the EP.

**Figure 3.**
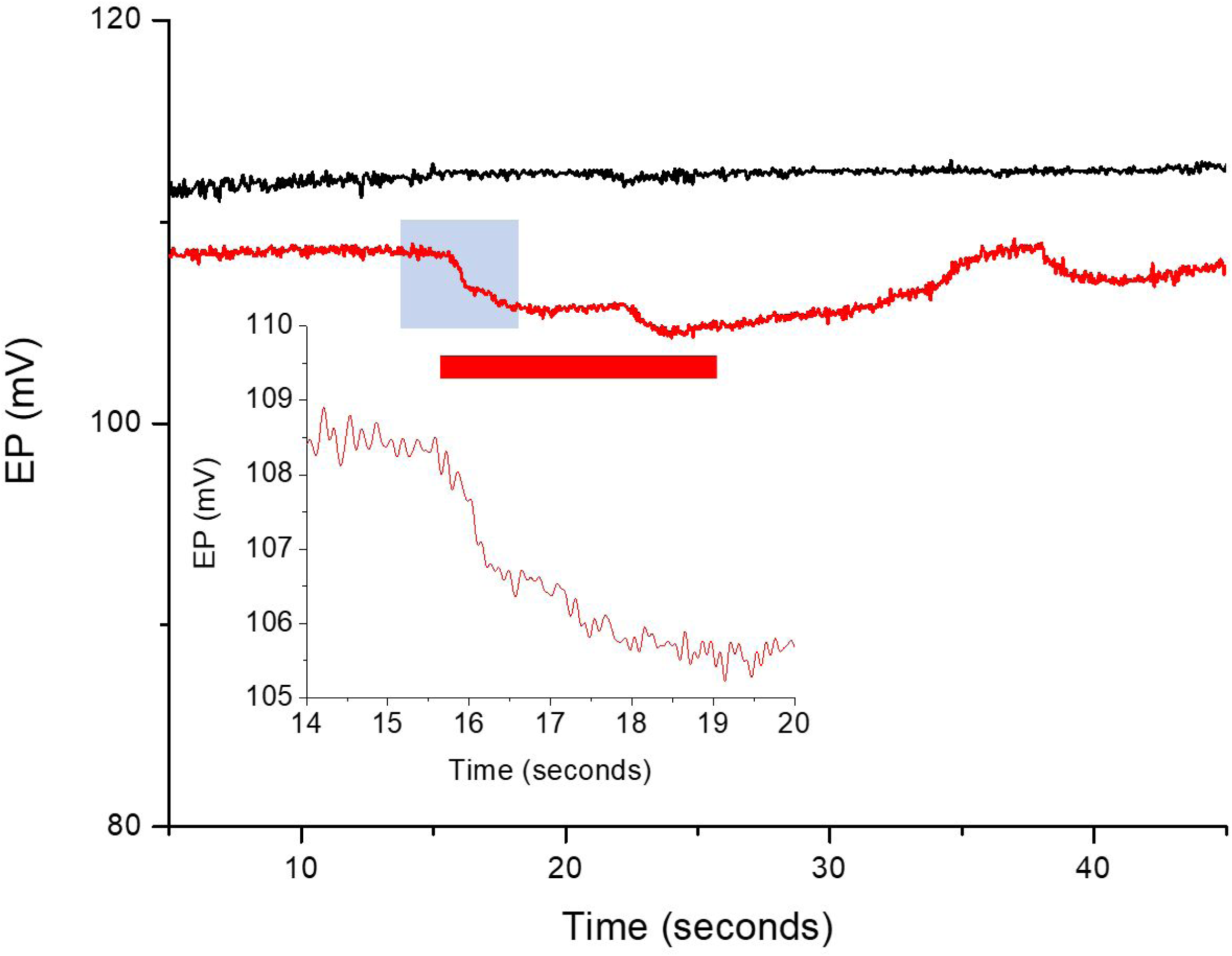
Prolonged activation of HOP causes small, prolonged reductions in the EP (red trace) recorded from scala media during a 10 s (red bar) illumination of the OoC. Black trace: EP from WT littermate that did not express HOP and did not respond to the OoC illumination. Inset: expanded EP taken from shaded area. Illuminating laser beam: 200 µm diameter, 0.25 mW*mm^-2^, 589 nm.

### HOP activation suppresses BM responses throughout their dynamic and frequency ranges

To examine how light activation of HOP expressing cochlear supporting cells influences tone-evoked BM mechanical responses, we measured the magnitude and phase of BM displacement as functions of SPL for different stimulus frequencies at, below, and above the CF of the measurement location, which was kept close to 51 kHz. BM frequency tuning curves are very similar to the frequency tuning curves of OHC electrical responses (Russell and Kössl, 1992a; Russell et al., 1995), and should provide a basis for comparing how OoC supporting cells influence OHCs and the mechanics of the cochlear partition when light activated.

We measured tone-evoked BM displacements with a self-mixing laser vibrometer (Lukashkin et al., 2005) in mice that expressed HOP in supporting cells and in WT littermates that lacked HOP expression. BM responses to tones were measured before, during and following a period of HOP activation by illuminating the BM in the immediate vicinity of the measurement site with the laser beam (200 µm diameter, 0.25 mW, mm^-2^, 589 nm). At all frequencies (34 to 60 kHz) and levels (20 to 100 dB SPL) that were measured, HOP excitation suppressed tone-evoked BM displacements resulting in a lateral shift of the level functions to higher SPLs (less sensitive, Figs. 4A-E) with a tendency to make the response-level relationship more linear. These reductions in sensitivity were usually associated with small phase lags of < 20° (Figs. 4A – E). Illumination of the BM had no observable influence on BM responses measured from WT mice (Fig. 4F). Due to desensitization (see Lukashkina et al., 2022) it was possible only to obtain one complete tuning curve (Fig. 4G) out of 10 preparations from which successful sensitive measurements had been made. Sensitivity at the 0.1 nm isoresponse thresholds of the level functions was reduced by 8.0 ±1.7 dB SPL, based on measurements from 15 level functions (n = 15), for frequencies 5 kHz above and below the CF and 14.2 ± 1.4 dB SPL, n =15, for frequencies at and close to CF. These sensitivity changes were associated with a phase lag measured at 70 dB SPL of 11.3 ±2.4 degrees, n = 15, throughout the frequency range of the tuning curve (Fig. 4G) as indicated in the sensitivity difference tuning curve (Fig. 4H). Suppression of BM displacement also caused a broadening of the frequency tuning as indicated by the Q_10 dB_ (Q_10 dB_ = CF divided by the bandwidth measured 10 dB from the tip), which was reduced from 8.6 to 3.6 without an obvious change in the CF. Thus, activation of HOP expressed in DCs suppresses tone evoked BM displacements, accompanied by small changes in phase, throughout their dynamic range and over a wide frequency range that includes levels and frequencies that are not subject to cochlear amplification at the recording location.

**Figure 4.**
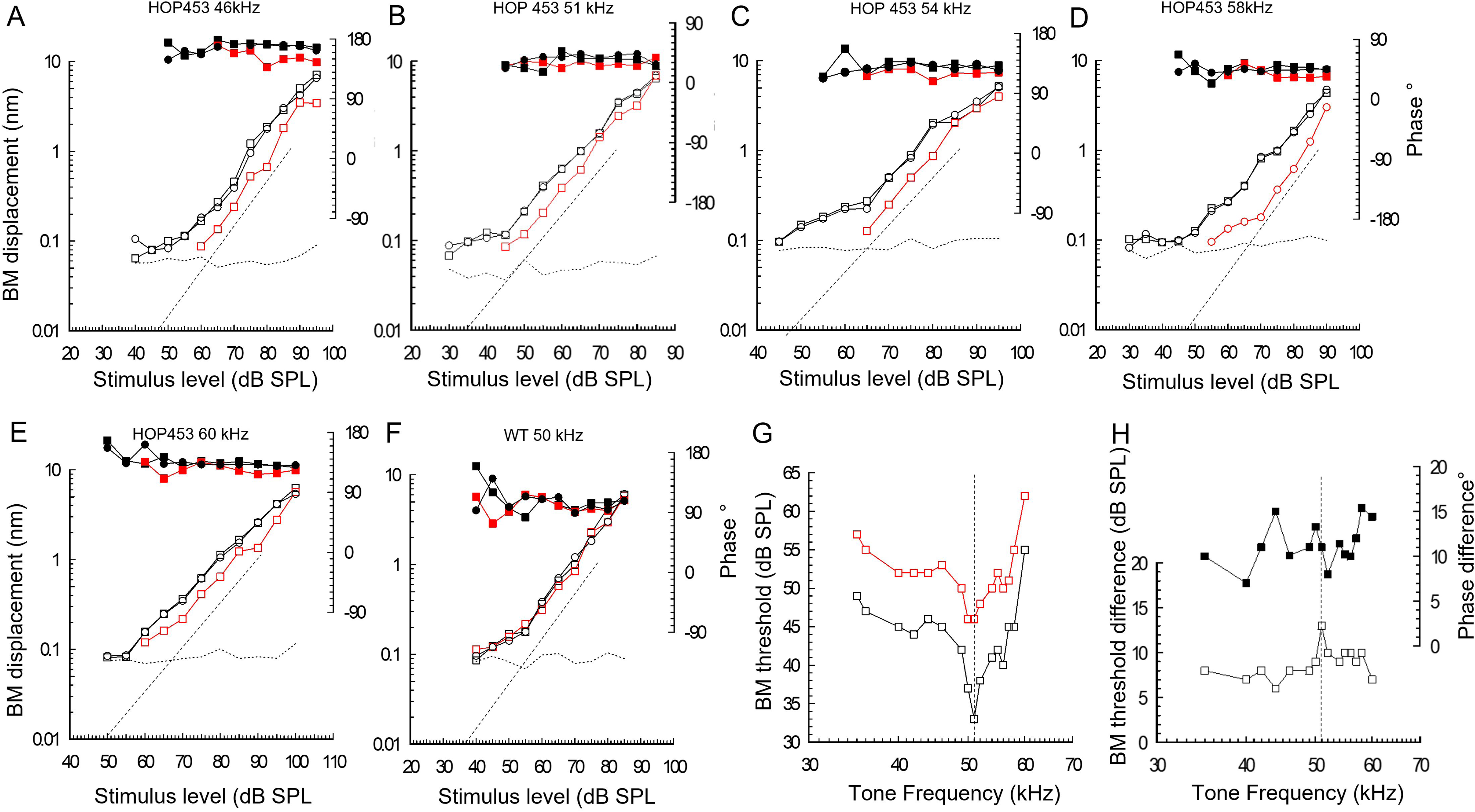
Light activation of HOP suppresses BM responses throughout their dynamic and frequency range. **A – F,** Magnitude (open symbols) and phase (solid symbols) of BM displacements to tones at the frequencies indicated as functions of sound pressure level (SPL) of the presented tones. Black squares: control before HOP light-activation. Red circles: during HOP light-activation. Black circles: following HOP light-activation. **A-E.** Measurements from HOP expressing cochlea. **F**. Measurements from a WT littermate that did not express HOP. **G**. BM displacement threshold (0.1 nm criterion) frequency tuning curve measured before (black squares) and during (red squares) light-activation of HOP expressing supporting cells. **E**. BM threshold difference between control and when HOP expressing supporting cells are activated, taken from **G** (open squares). BM phase difference measured at 70 dB SPL between control and activation of HOP expressing supporting cells. Dotted lines: **A-F**, slope of one; **G**, **H**, CF. Level function measurements were separated by periods of 5 minutes. Magnitude was corrected for middle ear transfer characteristics (Dong et al., 2013). Phase was measured relative to the speaker control voltage. Illuminating laser beam: 200 µm diameter, 0.25 mW. mm^-2^, 589 nm.

### HOP activation increases ERPDC negativity, alters ERP symmetry and inhibits CAP to low-frequency tones well below the CF

OHC ERPs (Fig. 5A) and OHC RPs (Fig. 5B) develop a fast, level-dependent DC (ERPDC and OHC DC) component that appears immediately at tone onset in response to moderate to loud tones (Figs. 5A, B, D). The ERPDC occurs in response to tones at all frequencies including those (e.g., 5 kHz, Figs. 5A, B) which should not be subject to cochlear amplification because they are considerably below the CF (50 – 60 kHz) of the recording site (Robles and Ruggero, 2001). ERPDCs can be negative at moderate sound levels but, at least for frequencies below the CF of the measurement location, the polarity is positive for high stimulus sound levels (Figs. 5A, D). The appearance of the ERPDC is not associated with measurable changes in the magnitude and phase of the ERP (Fig. 5E). HOP activation causes the ERPDC to be more negative at all levels of its manifestation and does not influence the amplitude and phase of ERPs that are not subject to cochlear amplification (Fig. 5E) and does not influence the ERPs and ERPDCs of mice that do not express HOP (Fig. 5C, control).

**Figure 5.**
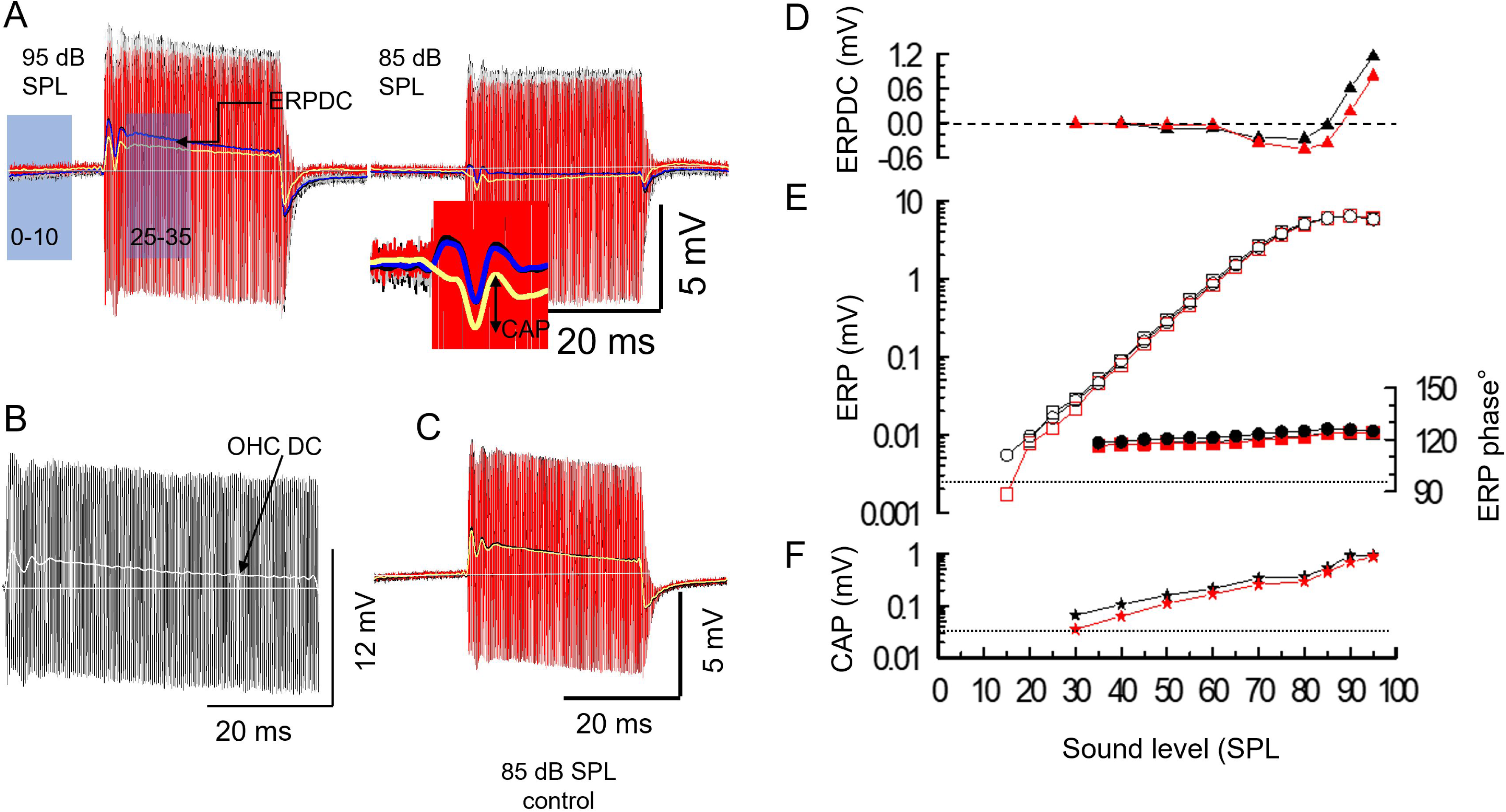
HOP activation increases ERPDC negativity, without altering ERP phase and magnitude, and attenuates compound action potentials to a low-frequency tone of 5 kHz. **A.** 85 and 95 dB SPL, 5 kHz ERPs from cochleae with HOP supporting cells. Black before, red during, grey following, BM illumination (to activate HOP). Superimposed lines: ERPDC (1 kHz FFT low pass filter of ERP). Black before, yellow during, blue following, HOP activation. White lines: CL potential. **B**. OHC intracellular receptor potential to 75 dB 5 kHz tones from WT mouse (resting membrane potential -42 mV). **C**. ERPs and ERPDCs from control mouse that did not express HOP. **D**. ERPDC amplitude (triangles) as function of SPL measured as the difference between measurements made in the 0 – 10 ms period before the tone burst and in the 25 – 35 ms period during the tone burst, as indicated in A, 95 dB SPL). Black triangles without HOP activation. Red triangles: during HOP activation. ERPDC measured as mean ± S.D of 2500 samples. ERPDC measurements with and without HOP activation were statistically very significantly different for levels > 50 dB SPL (two-tailed P value is less than 0.0001, t = 127.0260, df = 4998). **E.** Magnitude (open symbols), phase (closed symbols) of OHC ERP as functions of SPL for 5 kHz tones from a HOP mouse before (square), during (red) and following (circle) BM illumination of measurement site. Level function measurements separated by 5 minute intervals. Phase measured with respect to sound control voltage. **F**. CAP amplitude as functions of SPL, (stars, expanded insert, 85 dB SPL trace, **A**), before (black), during (red) HOP activation. Dashed line D: zero millivolts. Dotted lines E, F: measurement noise floor. Illuminating laser beam: 200 µm diameter, 0.25 mW. mm^-2^, 589 nm.

Sound processing in the OoC involves the transfer of sound-induced forces from the motile OHCs to the IHCs and the subsequent excitation of the auditory afferent nerve fibers (Robles and Ruggero, 2001). A remote measure of this process is the cochlear compound action potential (CAP), which can be measured from almost anywhere on the surface and within the cochlea itself, and at very low SPLs, can be used to monitor the frequency-dependent sensitivity of the cochlea (CAP audiogram, e.g. Russell and Sellick, 1977, Ozdamar and Dallos, 1978). With increasing SPL, the CAP becomes larger and wider due to the increasing contribution of afferent fibers from a broadening region of the OoC that is excited by the louder tones that extends from the cochlear frequency place to the most sensitive region of the cochlea (Lee et al., 2019). For example, IHC receptor potentials recorded from the 18 kHz CF region of the guinea pig cochlea can be elicited in response to tones at frequencies below 100 Hz at moderate to high SPLs (Russell and Sellick, 1983).

With these ideas in mind, we measured the amplitude of the CAP (see inset Fig, 5A, 85 dB SPL) as a function of SPL before and during HOP light-activation. At all levels, except at 95 dB SPL, the CAP was reduced in amplitude during HOP activation, reduction being greatest at low SPLs (Fig. 5F). CAP for tones below 95 dB SPL statistically significantly reduced during HOP activation over the level range 30 dB SPL to 90 dB SPL to 0.77 ± 0.05, n =8, of control, (two-tailed P value is equal to or less than 0.0004, t = 4.3853, df = 18, n = 10 for each measurement point). There was no measurable change in latency. An implication of this finding is that the CAP recorded in the CL is dominated by signals from afferent fibers in the immediate vicinity of the recording site and that their activity is modified by activation of HOP expressed in DCs located within the 200 µm of the beam. Furthermore, tone driven excitation of the fibers is reduced during HOP activation. CAP was reduced at all levels, including close to detection threshold, (∼ 80 µV at 30 dB SPL). HOP activation has no apparent influence on the amplitude and phase of ERPs, but it alters the magnitude and sign of the ERPDC. This could indicate that HOP activation alters the symmetry and operating point of OHC MET. Reduction in the CAP may indicate the transfer of forces between the OHCs and IHCs is reduced during HOP activation. The immediate appearance of the ERPDC and OHC DC receptor potential at tone onset could indicate that the asymmetry in DC component of the ERP and OHC receptor potentials is due to tonic radial shear displacement between the RL and TM which biases the operating point of the OHC MET in response to moderate to loud tones.

### HOP activation reversibly suppresses amplified ERPs, ERPDCs become more negative, and CAPs are reduced

ERP responses to tones that are subject to cochlear amplification (Fig. 6A, 44 kHz and Fig. 6B, 53 kHz) are reversibly suppressed for levels below ∼ 70 dB SPL when HOP-expressing supporting cells are light activated (Figs. 6C, D). Under these circumstances, illumination of the HOP-expressing supporting cells reduces the magnitude of the ERPs equivalent to a change in gain of 13.5 ± 2.7 dB, n = 7 at 25 dB SPL when the tone is at the CF of 53 kHz of the recording site (Fig. 6D). The reduction in the ERP is smaller at frequencies below the CF (4.5 ± 1.6 dB, n = 8 at 40 dB SPL, tone = 44 kHz, Fig. 6D). Suppression is due largely to increased linearity (dashed lines, Figs. 6C, D) of the slope of the ERP level function with HOP activation and is progressively reduced with increasing level. Associated with the suppression of responses that can be subject to cochlear amplification is a small phase lag of less than 20 degrees. Illumination of the BM that caused suppression of the ERP and increased negativity of the ERPDC (see below) in mice with HOP-expressing supporting cells, had no observable influence on ERPs and ERPDCs in WT mice that did not express HOP in the supporting cells (Fig. 6E).

**Figure 6.**
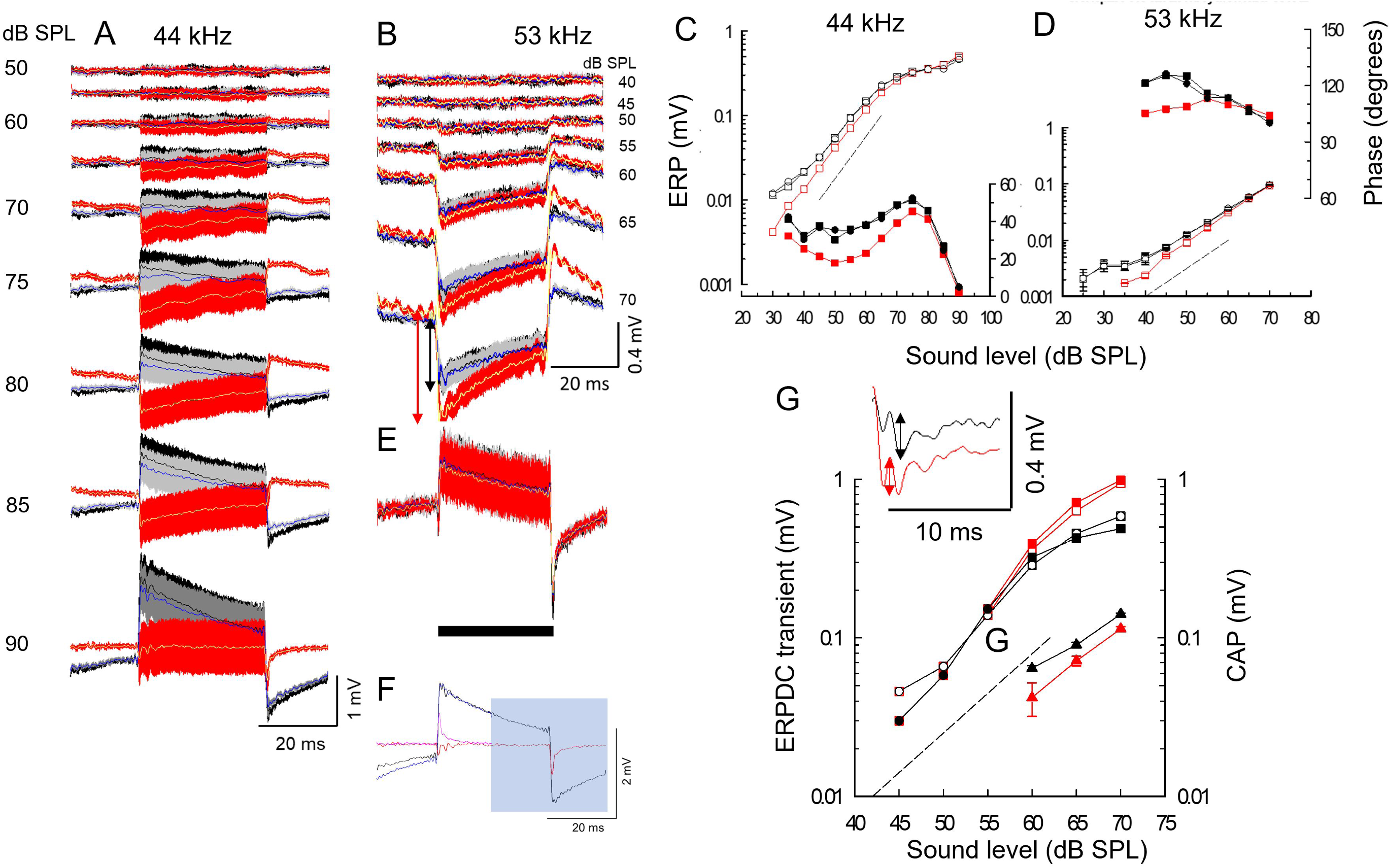
HOP activation reversibly suppresses amplified ERPs, ERPDCs become more negative and CAPs are reduced. **A**. ERPs in response to 50 – 90 dB SPL 44 kHz tones. Before (black), during (red), after (grey), illumination of BM at the measurement site. Superimposed fine traces: ERPDC (FFT low pass filtered (1 kHz) raw ERP). Before (black), during (yellow), after (blue) BM illumination. **B**. Similar to **A** but CF (53 kHz) of recording location. **C, D**. Magnitude (**A** ± S.D, n = 5 successive measurements) and phase of 44 kHz (**C**) and 53 kHz (**D**) ERPs tones as functions of SPL before (black open square), during (red open square) and following (black open circle) BM illumination. Phase: solid symbols. Level function measurements separated by 5 minute periods. **E**. ERPs and ERPDCs, 44 kHz, 90 dB SPL, WT mouse without HOP expression. **F.** ERPDC before, black trace and during HOP activation, red trace from **A**, 90 dB SPL. Offset transient, blue shaded flipped about the x and y axes and replotted over the onset transient. **G.** Amplitudes of ERPDC onset (open symbols) and offset (closed symbols) transients (arrows **B,** 70) as functions of SPL with signs of onset responses changed to positive to enable comparison of onset and offset responses (black before, red during BM illumination). Triangles, magnitude of CAP responses to 53 kHz tones (mean ± SD, n = 5) as functions of SPL (see inset CAP, 70 dB SPL 53 kHz tones, double headed arrows). Black triangles, before; red triangles, during BM illumination. ERP measurements uncompensated for microelectrode time constant. 44 kHz traces: FFT band blocked (1 - 40 kHz), 53 kHz traces: (1 - 50 kHz). Illuminating laser beam: 200 µm diameter, 0.25 mW. mm^-2^, 589 nm. FFT (fast Fourier Transform).

ERPs generated in response to tones at frequencies a half-octave below the CF undergo a sharp phase lag when the tone level exceeds about 80 dB SPL, as described previously for the gerbil (Zwislocki, 1988) and guinea pig cochleae (Russell and Kössl, 1992a, b). OHCs are excited by displacement of the stereocilia bundle toward the tallest row (Russell et al., 1986). According to a mechanical model of the cochlear partition (Zwislocki, 1986), this can be achieved by displacements of the BM towards either the scala vestibuli or the scala tympani, depending on the relationship between the rotational stiffness of the OHC stereocilia bundle and the TM impedance, which should be minimal at the TM resonant frequency (Allen, 1980; Zwislocki, 1986; Kössl and Russell, 1992 Gummer et al., 1998; Lukashkin et al., 2010; Nankali et al., 2022; Lukashkin et al., 2023). A level-dependent change in this relationship, possibly as a consequence of level-dependent changes in the mechanical impedance of elements within the cochlear partition, could reverse the direction of excitation (Zwislocki, 1986, 1988; Lukashkin et al., 2010). HOP activation, however, does not influence this phase reversal of OHC excitation or indeed, significantly influence the phase of the ERP for levels above ∼75 dB SPL, which is when the ERP phase change begins and OHC responses are no longer amplified (Fig. 6D).

ERPDCs generated in response to CF tones (53 kHz) are detectable above the measurement noise floor for levels above ∼ 45 dB SPL (Figs. 6B, G) and from levels above ∼ 55 dB SPL in responses to tones below the CF (44 kHz, Figs. 6A, F), which is above the frequency (38 kHz, a half-octave below CF) where responses are not subject to cochlear amplification (Robles and Ruggero, 2001). They appear at levels when the contribution of cochlear amplification to the OHC voltage responses and the ERP become disappearingly small and when HOP activation has little or no influence on the magnitude and phase of the ERP (Fig. 7C, D). The ERPDC first appears either as a positive potential for tones at frequencies below the CF, e.g., at 44 kHz (Fig. 6A), or negative e.g., at the 53 kHz CF (Fig. 6B) and it remains almost constant in amplitude for the duration of the tone and falls immediately to the CL potential at tone offset. With increasing SPLs, the ERPDC increases in level but is accompanied by an additional component that attempts to restore the DC to the CL potential (0 mV) for the duration of the tone (see also Figs. 5, A, C). For positive ERPDC this results in a drift of the DC in the negative direction and in the positive direction for negative ERPDC. At tone offset, there is an ERPDC offset transient equal in amplitude, but of opposite in sign to the ERPDC onset transient, that declines over time to the level of the CL potential. Restoration of the ERPDC to the baseline at tone offset has the same magnitude and time dependency as the restoration that takes place during the tone burst, but of the opposite sign. This point is illustrated by flipping both dimensions of the offset traces shaded in blue and fitting them to the onset transients (Fig. 6F). The rate of restoration (not explored here) appears to depend on the level of the ERPDC and declines faster when the deviation of the ERPDC is greatest, regardless of the polarity of the ERPDC. When the same sequence of acoustic stimulation is performed together with HOP activation, the ERPDC becomes more negative, thereby implicating roles for the DCs and OPCs in the generation of the ERPDC. The relationships between the onset magnitude of the ERPDC as a function of the level of the 53 kHz tone (Figs. 6B) are shown in Fig. 6G. The positive ERPDC increases as a function of the of the tone level (Fig.6, black symbols) as does the negativity caused by HOP activation (Fig 6G, solid red symbols).

Over a measurement range (60 - 70 dB SPL) where CAPs could be measured unambiguously in five cochleae, HOP activation suppresses the amplitude of CAPs recorded in the CL (Fig. 6G), which provided indirect evidence that transmission of forces from the OHCs to the IHCs is reduced. It is evident that through activating HOP at frequencies when the control of cochlear amplification is dominated by the ERP, DCs and OPCs have roles in optimizing cochlear amplification at low stimulus levels. At all frequencies and levels explored in the measurements reported here, DCs and OPCs optimize the transmission of OHC forces to IHCs and control the OHC MET operating point.

## DISCUSSION

Activation of HOP expressed in the cell membranes of isolated DCs caused them to be swiftly and tonically hyperpolarized while HOP activation of DCs and OPCs *in vivo* caused a slow, irregular reduction in EP likely involving changes in the gap junction network that interconnects DCs. Such changes could potentially influence the mechanics, electrophysiology, and electrochemistry of the OoC (Jagger and Forge, 2015; Mammano, 2019; Lukashkina et al., 2022; Levic et al., 2022). HOP activation suppressed cochlear amplification, revealed roles for DCs and OPCs in setting the OHC MET operating point and in transfer of sound-induced forces from OHCs to IHCs and afferent neural excitation. HOP activation suppressed BM responses to tones at all levels including at frequencies that do not receive cochlear amplification and revealed DCs and OPCs control forces transmitted between the OoC and OHCs transverse, radial and longitudinal directions.

### HOP activation suppresses cochlear amplification and modulates ERPDCs

HOP activation causes hyperpolarization of DCs and OPCs through the inward transport of Cl^-1^ from the OoC Cortilymph leading to suppression of cochlear voltage responses to tones and levels that are subject to cochlear amplification (frequencies above a half-octave below the CF and levels below ∼70 dB SPL, Robles and Ruggero, 2001) and suppression of tone-evoked BM displacements including those at levels and frequencies not subject to amplification.

Suppression of cochlear amplification could be due to one or a combination of processes. Cl^-1^ ions interact with the OHC motor protein prestin to confer its voltage-dependent motility (reviewed by Ashmore, 2008; 2021). Cortilymph ionic composition is rapidly changed by the activity of the cells that enclose it (Johnstone et al, 1989) and HOP activation might be expected to reduce Cortilymph Cl^-1^ levels. Reduction in perilymph Cl^-1^ levels reduces amplification of BM displacement and, presumably, OHC voltage responses (Santos Sacchi et al., 2006). Another possibility is based on the calcium-dependent motility of DCs and OPCs (Dulon et al.,1994; Vélez-Ortega et al, 2023) and finding that voltage changes in DCs can directly modulate OHC electromotility through DC-OHC mechanical coupling (Yu and Zhao, 2009), perhaps by altering OHC turgor pressure to which prestin motility is sensitive (Kakehata and Santos-Sacchi, 1995). Either mechanism working alone or in tandem would attenuate cochlear amplification rather than setting it in a resting mode by disabling fine control of cochlear amplification as occurs through activation of ChR2 (Lukashkina et al., 2022). HOP activation always caused a phase lag in sound-induced mechanical and voltage responses, whereas ChR2 activation caused both phase lags and advances relative to the control. The lags could indicate a further basis for reduced cochlear amplification. Models and *in vivo* measurements reveal OHC force delivery at maximum BM velocity is important for optimal cochlear amplification (Geisler, and Sang, 1995; Nobili and Mammano, 1996; Russell and Nilsen, 1997).

At levels and frequencies where OHC prestin-based motility plays little or no role in shaping cochlear responses, we found that HOP activation attenuates BM responses to tones and modifies the tonic ERPDC, but not the phasic ERP. We suggest that the changes in OoC mechanical responses and the appearance of the ERPDC may be due to the consequences of passive mechanical interaction between the OHCs and their supporting cell scaffolding. During intense sound stimulation, DCs and Hensen’s cells reversibly change shape and structurally modify the organization of the OoC (Flock et al., 1999) and TRPA1, that is expressed in Hensen’s cells and could be activated during acoustic trauma, causes prolonged Ca^2+^ responses, which propagate across the OoC and cause long-lasting contractions of the motile pillar and Deiters’ cells (Vélez-Ortega et al., 2023). This proposed interaction could stimulate stretch activated conductances that populate OHC lateral membranes (Iwasa, et al., 1991; Ding et al., 1991; Rybalchenko and Santos-Sacchi, 2003; Wu et al., 2017) resulting in sinusoidal OHC length changes orthogonal to the stimulus direction together with a tonic increase in OHC length that depends on the amplitude of the near field component of the acoustic-mechanical stimulus (Brundin and Russell, 1993). At low-stimulus levels, OHC tonic displacements do not adapt throughout stimulus duration. With increasing levels, OHC tonic displacements adapt, as if caused by a relaxation in OHC axial stiffness. At stimulus offset, OHC length relaxes to its resting length with a time course similar to that of the on-stimulus relaxation. The proposed relaxation in axial stiffness observed during mechanically induced elongation of isolated OHCs could be the basis of the apparent adaptation of the ERPDC that occurs during moderate to high level tones and following tone offset.

### HOP Activation impairs force transmission between outer and inner hair cells

IHC hair bundles are displaced through sound-induced radial shear displacements between the TM and reticular laminar. *In vivo* intracellular measurements of IHC low-frequency receptor potentials in the mid to high frequency turns of the guinea pig cochlea (Sellick and Russell, 1980; Nuttall et al., 1981; Russell and Sellick, 1983; Dallos, 1986; Patuzzi and Yates, 1987) are consistent with a proposal (Dallos et al, 1972) that IHC hair bundle displacement is related to BM velocity at low frequencies. Changes in the spatial relationship between the TM and reticular laminar, relative angles between TM and RL, and radial displacements between the TM and RL, all of which could involve the OHC cellular scaffolding, can influence the fluid coupling of the IHCs to shear between the TM and RL (Nowotny and Gummer, 2006; Prodanovic et al., 2015; He et al., 2018; Sasmal and Grosh, 2019). With respect to this, HOP activation during low-frequency, high-level tone stimulation increased the negativity of the ERPDC without changing the magnitude and phase of the ERP. However, the symmetry and operating point of the ERP, and hence that of the OHCs, was changed away from its most sensitive operating point (Russell et al., 1986; Russell and Kössl, 1992b) which, together with HOP-induced changes to the OHC cellular scaffolding and IHC fluid coupling, reduced force transfer between the OHCs and IHCs thereby reducing afferent excitation and hence the magnitude of the CAP.

### HOP activation implicates roles for DCs and OPCs in transfer of longitudinal cochlear forces

HOP activation desensitizes BM threshold displacement tuning curves by 10-15 dB SPL over the entire measured frequency range with maximum suppression and broadening at the tuning curve tip where tone-evoked BM responses are subject to greatest cochlear amplification. It seems, therefore, that HOP activation causes mechanical changes in the DCs and OPCs that alter both the active and passive mechanical properties of the cochlear partition. Because the supporting cells, including DCs, are thought to shape energy transmission along the cochlea (Geisler and Sang,1995; Russell and Nilson, 1997; Steele et al., 1999; Liu et al., 2022), the observed changes in responses due to HOP activation might be not only due to local mechanical changes at the CF location but because of mechanical changes basal to the location. In this respect, HOP induced suppression of BM responses contrasts with inhibition directed specifically at the OHCs through stimulation of their efferent innervation which causes maximum reduction of BM displacement responses to CF tones, where the contribution of cochlear amplification is greatest, and little or no inhibition of BM displacements in responses to stimulus frequencies that are not subject to cochlear amplification (Murugasu and Russell, 1996; Cooper and Guinan, 2003). However, HOP activation suppresses BM responses over a broad frequency range of at least 35 kHz to 62 kHz. It is suggested that OPC and DC activation in the immediate beam of the laser suppress all movements of the BM regardless of their frequency, amplitude, and origin. This might include movements of the cochlear partition originating from frequency regions basal to the CF where HOP was activated and the force transmission of DCs and OPCs were affected. Movement from remote sources could reach the measurement site on the BM through longitudinal transmission of forces along the cochlear partition, which might implicate a role for supporting cells, including DCs and OPCs as proposed in the “feedforward” models (Geisler and Sang,1995; Russell and Nilson, 1997; Steele et al., 1999; Liu et al., 2022). These forces do not appear to excite OHCs, which respond to radial shear between the TM and RL and have tuning curves resembling the BM frequency tuning curve (Russell and Kössl, 1992; Russell et al., 1995) and do not have the broad frequency characteristics reported for the frequency tuning of the RL (He et al., 2018; Ren et al., 2016) and the structures contributing to the frequency characteristics of the “hotspot” within the OoC (Cooper et al., 2018) which may be generated by longitudinal flow of forces along the cochlear partition length.

The findings, reported here, that hyperpolarizing HOP activation of DCs and OPCs reduces cochlear amplification, reduces the movement of the BM at all levels and frequencies, and reduces high level IHC excitation, and depolarizing COP activation reduces the recovery time from intense sound stimulation (Lukashkina et al. 2022), indicates DCs and OPCs as potential targets for investigating the prevention and treatment of noise induced hearing loss.

## Acknowledgements

The authors thank Sarath Vijayakumar and Cassidy Nguyen for technical assistance on mouse colony management at Creighton University.

## Author contributions

IJR, ANL and JZ, conceived and designed the study. JZ designed mice with HOP expressing cochlear supporting cells. JAD designed HOP mouse cross experiments and contributed to genotyping. SL designed *ex vivo* measurements. ANL and IJR designed *in vivo* measurements. PS and SL measured and analyzed data from *ex vivo* experiments. VAL and IJR measured and analyzed data from *in vivo* experiments. SL prepared cochleae for immunohistochemistry and ZX performed cochlear immunohistochemistry. VAL organized mouse breeding at Brighton. ANL wrote computer programs. IJR and ANL wrote the paper with contributions from the other authors.

## Data and materials availability

All data is available in the main text. Additional data related to this paper may be requested from the authors.

